# Ecoduality

**DOI:** 10.1101/2025.06.21.660866

**Authors:** Robert E. Ulanowicz

**Affiliations:** University of Maryland Center for Environmental Science; University of Florida Arthur R Marshall Laboratory for Theoretical Ecology

## Abstract

Dualism has been a viable concept in physics since the 1920s, when Arthur Compton demonstrated the corpuscular nature of photons as distinct from its wave nature. More recently, information theory applied to ecological dynamics has portrayed the behavior of whole ecosystems as being, not only dual, but actually dialectical (oppositional) in nature. Duality in ecology, however, pertains not only to whole systems but also to the constitutive elements as well – their taxa and exchanges. Here the quantitative degrees to which each element contributes antagonistically to system structure and flexibility is parsed out. The analysis reveals a more comprehensive alternative to the use of Boltzmann-Gibbs “entropy” with which to express biodiversity. Virtually all real ecosystem elements exhibit such quantifiable duality, which depend more on system topology than on positivist mechanisms. One is led, therefore, to search among realms beyond conventional physics, such as process theory and apophasis (lacunae) to formulate a complete quantitative description of ecosystem status and behavior. Because numerous other complex systems can be described in terms of two-dimensional flow matrices, the possibility arises that dualistic behavior may be far more widespread in nature than heretofore has been intuited.

---

*“The test of a first-rate intelligence is the ability to hold two opposing ideas in mind at the same time and still retain the ability to function*.*”*

F. Scott Fitzgerald, *The Crack-Up*.

## Introduction

Since about 1940, conventional ecosystems theory, like physics, has dealt largely with objects and mechanisms, so that ecosystem behavior could be described in 2009 by Francisco Ayala as merely “objects moving according to universal laws”^1^. This conclusion was drawn despite the experimental discovery in the 1920s that photons could exhibit the dual natures of mass and waves^2^, which still poses a logical paradox. Classical physics remains inadequate to support such dualistic behavior. Could there be something missing from contemporary physics that might support dialectical phenomena that function antagonistically?

One might begin by noting that the universal force laws are capable of dealing with only homogeneous attributes^3^, whereas ecosystems, by definition, encompass seriously heterogeneous collections of taxa and flows. Such extreme variation renders the boundary constraints on realistic ecosystems exponentially combinatoric, thereby defeating any formulation in closed form^4^. The failure of homogeneous laws to describe fully the behavior of massively heterogeneous systems prompts an alternative approach, such as one that casts complex systems as networks of elements connected by flows of material, energy or information.

It bears mention as well, that photons and electrons are among the simplest of all objects in the cosmos. Might more complex systems be capable of such dualistic ontology, *a-fortiori*?

An advantage of regarding systems as networks allows that they can exhibit valuable insights into system without regard to their constitutive eliciting causalities. Conventionally, flows are considered as resulting from causes arising within the elements they affect. However, configurations of processes may also combine to create new taxa^5^. Instead of invoking differential equations (which require causal attributions), entire networks can be analyzed phenomenologically (i.e., absent causes) to describe the quantitative behavior and development at the system level^6^. In fact, applying information theory to network structures has already revealed a dialectical opposition between functions that constrain the behaviors of the network nodes and those that quantify the residual independence of the same elements^7^. Such a revelation arose out of an approach that has been called “process ecology^8^“.

In order to treat the dialectical nature of ecosystem dynamics, it is important to focus on an important characteristic of the system called “entropy”^9^. The second law of thermodynamics, expressed in statistical terms, requires that system disorder at universal scales always increases. That this law remains inviolate, leads many to believe that increasing entropy is the final cause behind all behavior in the universe. But such attitude seriously exaggerates the power of entropy. The formulation quantifying entropy by Ludwig von Boltzmann and Josiah Willard Gibbs was predicated upon the *absence* of any constraints among system elements (the definition of an ideal gas.) That is, entropy, or the lack of any constraint, is identical to an *apophasis*, a name given to something that *does not exist*. This is in contrast to the backbone of physics, which consists of positivist features, such as mass, energy, momentum etc., which are capable of causing events to happen. Entropy, instead, is like a vacuum that passively *allows* room for any adjacent cotemporaneous positive causes to actively *make* things occur.^10^ The scenario is akin to the Asian interaction of Yin and Yan^11^.

But how can the role of apophasis be followed quantitatively and with data? How can the nonexistent be quantified at all? Actually, measuring that which is missing is no major stumbling block. It is done all the time. For example, “The glass is half full.” describes how much is missing in terms of the capacity of the glass. This is fully consistent with the classical laws of thermodynamics, because calculations of entropy are always made between two different states, which is to say entropy cannot be defined and measured without reference to some positivist absolute (a statement equivalent to the third law of thermodynamics.)

### Quantitative Considerations

The measure of the *capacity* of a system of *n* entities for action was conveniently and elegantly formulated, first by Boltzmann, and refined later by Gibbs (hereinafter referred to as the B-G formula.) They inconveniently identified the subject as the *entropy* of a system. The inconvenience arises because the formula was predicated upon the unrealistic assumption that the elements of the system were incapable of affecting one another – the definition of a so-called *ideal* gas. (No such gas exists in reality.) Because the convention in physics was upon objects, Gibbs’ formulation of Boltzmann’s law was promulgated as:

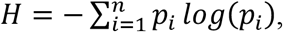

where *H* is the magnitude of system entropy and *p*_*i*_ is the relative density of objects in class *i*.

The inadequacy of *H* (like that of the ideal gas) was that it did not include the more important phenomenon of the interactions between the elements of the system. Fortunately, this lacuna is easily remedied by focusing instead on the *relationships* among the atoms. These can be represented by the joint probabilities of the interaction of any *i*^*th*^ element of the system with an arbitrary *j*^*th*^ entity, and will be denoted as *p*_*ij*_. Whence, the extended B-G measure of the possible Capacity for *Interaction*, C, within the system can be written as,

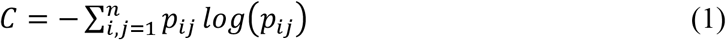

The marginal probability that the *i*^*th*^ element is involved becomes 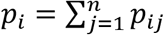 and for the *j*^*th*^ element 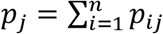. These emendations make it possible to decompose *C* into two separate groups of terms^12^ as,

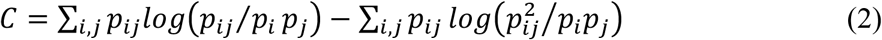

This decomposition tells us a lot more about how the system is behaving than does the aggregated B-G formula. (Unfortunately, it also requires more data to instantiate.) The first summation on the right-hand-side is called the Average Mutual Information (AMI). It should be noted that all the various forms of information share the nature of constraint, so that the AMI becomes a measure of the amount of system activity that is engaged in maintaining the kinetic form of the interaction scheme. One might think of it as gauging the aggregate formal cause (*sensu* Aristotle) that maintains the integrity of the system.

The second group of terms is known in information theory as the conditional entropy (Φ) of the ensemble. In contrast to the AMI, it portrays the relative independence and flexibility of the network of actions. One aspect of *Φ* is that it represents the strength in reserve that the system retains to reconfigure itself to adapt to novel perturbations. In abbreviated form,

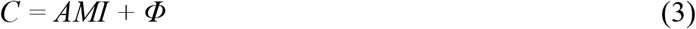

Both of these partitions can be shown to be non-negative (*AMI* ≥ 0; *Φ* ≥ 0.)^13^ This implies that a change in any probability will elicit a complementary change between the two groups. In other words, an alteration in either sum will induce an opposing change in the other. This quantitative expansion of the system capacity reveals a true Hegelian *dialectic* between constraint or form vs. freedom or flexibility.^14^ Although the characteristics are opposing, they are mutually obligate -- the first for performance and the second for persistence.

### A Dialectical Worldview

When the capacities of numerous ecological systems were expressed in terms of *AMI* and *Φ*, the ratios between them tended to cluster around 40% *AMI* and 60% Φ. When the same exercise was carried out on economic cash flow networks they tended to cluster at lower levels of constraint (*AMI/C*), between 15 and 20%^15^. Full networks of entire organisms are more difficult to estimate, but intuition would suggest that they would be even more ordered than ecosystems.

Still, all three classes of systems can be portrayed and analyzed to reveal a balance in the tension between constraint and flexibility (*AMI* vs *Φ*). Because all living systems can be represented as complex networks, it follows that the dynamics of living systems is more than simply “objects moving according to universal laws.”

Living systems, then, by virtue of their complex nature, exhibit dualistic behavior, but that dualism is not causally symmetric. As discussed above, constraints (*AMI*) are maintained by positivist attributes, such as force, mass, momentum, information, etc. These are the factors that generically “*make* things happen.” In contrast, the entropic portion of system capacity (*Φ*) expresses that which does not exist (order) or which no longer exists (the absentia or apophatic). It does not actively make events happen, but rather *allows* action by the positivist features. Real changes require both features of capacity, that is, active *and* passive, to arise and persist.

### Individual Contributions to System Dynamics

One can go further. Examination of equation (2) reveals that each interaction between *i* and *j* with joint probability *p*_*ij*_ modulates one and only one term in the sums that define *AMI* and *Φ*. That is, each individual term consists of a *p*_*ij*_ that multiplies (scales) a logarithmic function of the other actors in the network. In simple terms, each interaction modulates the contribution that the *i-j* relationship makes, respectively, to the active and passive dynamics of the whole system. One can thus quantitatively assess the relative role that each interaction contributes to either the active (*AMI*) or passive (*Φ*) nature of the system.

To assess these contributions, one needs to compare each *i-j* term in their definitions. That is, one needs compare *AMI*_*ij*_ with *Φ*_*ij*_, where,

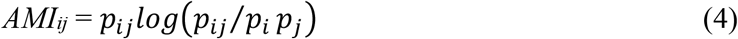

and

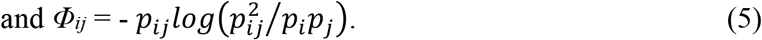

In functional terms, *Φ*_*ij*_ gauges how much the *i – j*^*th*^ relationship augments system freedom and flexibility (lack of constraint), allowing it to adapt to perturbations. It is easy to show that each *Φ*_*ij*_ is inherently non-negative: As is standard practice in statistics, all probabilities are assumed to be normalized over the closed interval [0,1]. The marginal sums, *p*_*i*_ and *p*_*j*_ both include the common joint probability in their sums and thus are both ≥ *p*_*ij*_. Whence, 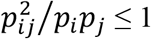, making its logarithm inherently non-positive, and the negative sign that precedes the logarithm ensures that the contribution *Φ*_*ij*_ is non-negative. That is, any *i - j* interaction in any network (other than an equiponderant straight chain) will make a positive contribution to the system’s conditional entropy.

The influence of the positivist constraint that relationship *i - j* exerts upon the kinetic form of the whole system is measured by *AMI*_*ij*_. Ordering the links in descending order of their *AMI*_*ij*_s reveals which flows are most significant in defining current system structure. Unlike the *Φ*_*ij*_, however, there is no guarantee that every *AMI*_*ij*_ will be ≥ 0. Some, on occasion, can be negative, which indicates that the two nodes occur less frequently together than would be expected if they were independent. In other words, the link, *p*_*ij*_, departs from its equilibrium value, *p*_*i*_*p*_*j*_. in that the pair members are mutually exclusive to an extent. Like the positive *AMI*_*ij*_, that binds the pairs, partial mutual exclusivity (a negative *AMI*_*ij*_) also serves as a structural constraint affecting system form.

It is important to note again that each relationship evinces antagonistic tendencies. Unlike with particle duality, which simply states that the same entity behaves differently in different contexts, network duality portrays inherent contradictory tendencies towards constraint and/or apophasis. It follows that their causal natures are distinct – positivistic effects that serve to *make* or preserve form vs. apophatic allowance that *permits* changes to occur.

### Quantifying Duality of Function in Real Ecosystem Networks

So how might these metrics work when expressed in terms of real ecosystem data? First it would be helpful to choose a network that has appeared in publication and has appropriate dimension. That is, sample networks that are too simple often do not satisfactorily portray system properties. Bersier & Sugihara^16^, for example, observed that communities with fewer than 12 members behaved qualitatively differently and sometimes spuriously than more fully articulated ones. On the other hand, systems with numerous nodes become too complicated to illustrate graphically. A simple intermediary example is the 16-member trophic network of carbon flows in the ecosystem of Crystal River, Florida (Figure 1).

**Figure 1.**
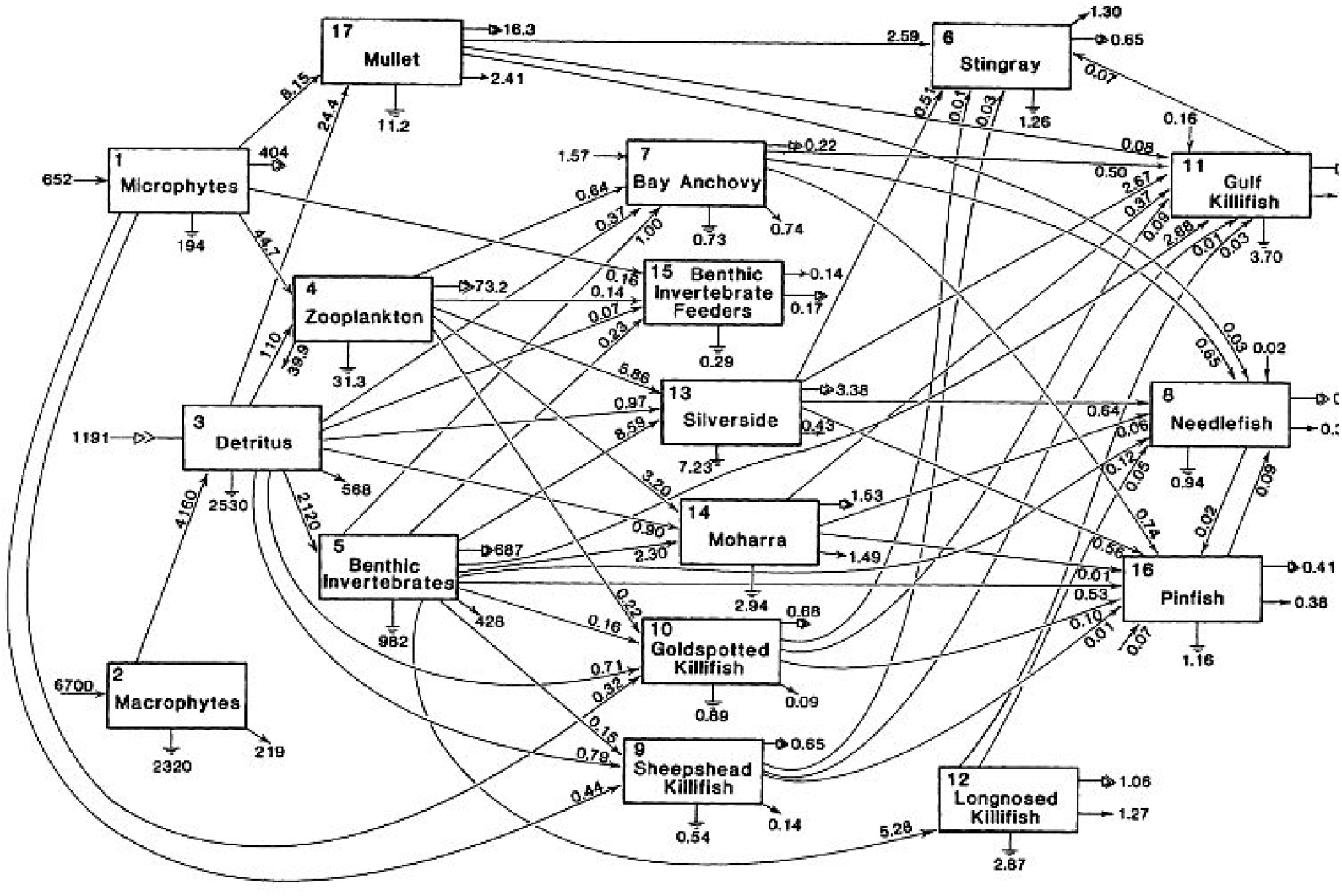
A diagram of the fish community in a Crystal River, Florida tidal marsh gut (M. Homer and M Kemp, unpublished.) The magnitudes of all flows are given in mgC/m^2^-y. Exogenous inputs are arrows without tails, while respirational dissipations are reported under the ground symbols, and exogenous exports are denoted by simple arrows not ending in a compartment. Compound arrowheads designate material recycled to detritus.

But even this simple ecological example consists of 91 flows and becomes cumbersome as an exercise for publication. Those wishing to examine pertinent results from the full analysis of the Crystal River Marsh system and for a 125-node representation of an Everglades Cypress Marsh network can request the full results from the author

To preserve space and ink, most of this presentation will be centered on an early 5-node representation of energy transfer in the Cone Spring system in Iowa (Figure 2).^17^ What it lacks in ecological reality is compensated for by its analytical simplicity. It consists of only 16 exchanges.

**Figure 2.**
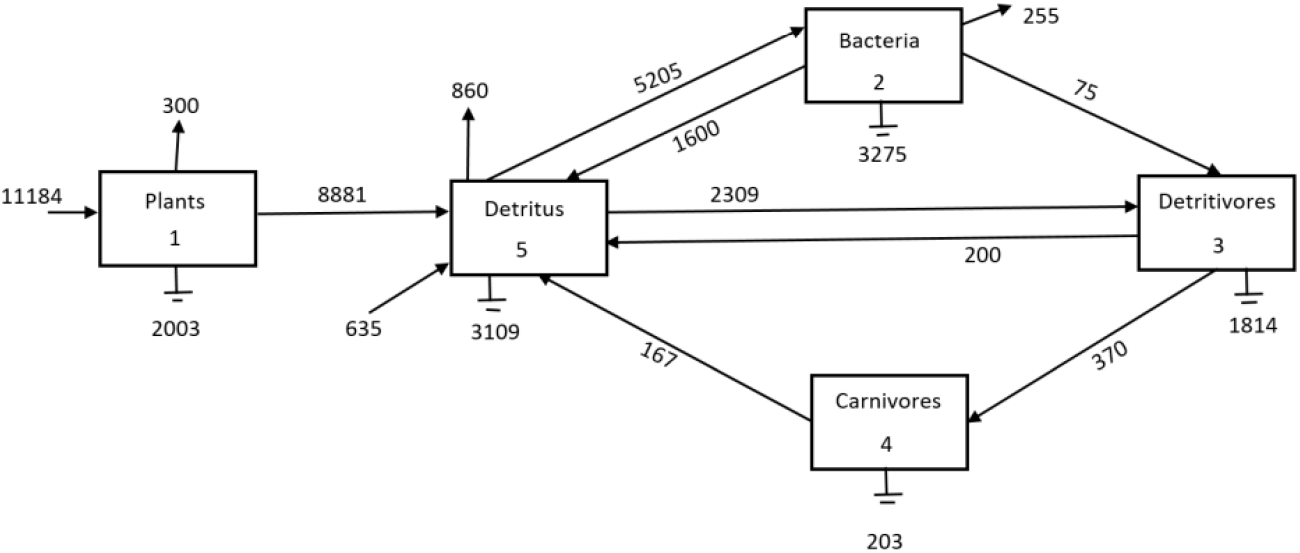
Energy exchanges in Cone Springs

The Cone Spring network (Figure2) can be enumerated as a 7-dimensional matrix of exchanges (Table 1). The *i – j*^*th*^ entry is the amount of embodied energy (in kcal/m^2^-y) that flows from the donor taxon in the *i*^*th*^ row to the recipient in the *j*^*th*^ column. Inputs from outside the system into the *j*^*th*^ taxon appear in the *j*^*th*^ entry of row 6, while exports from the *i*^*th*^ species is denoted by its respective entry in column 6.^18^ This array makes it easy to estimate all the probabilities that appear in Equation (2). The marginal sums of each column *j* appear in the *j*^*th*^ entry of row 7, whereas the marginal sums of the donations of the *i*^*th*^ taxon are placed in their respective element of column 7. The system was estimated to be in seasonal steady-state so that the sum of all marginal inputs denoted in row 7 is equal to the sum of all marginal donations tallied in column 7 (not a necessary restriction.) That value is the sum of all flows in the network, which by convention is denoted as the Total System Throughput (TST) in row 7, column 7.

**Table 1.** Flows (kcal/m^2^-y) and their marginal sums in Cone Spring.

|  | 1 | 2 | 3 | 4 | 5 | 6 | 7 |
| --- | --- | --- | --- | --- | --- | --- | --- |
| 1 | .0 | .0 | .0 | .0 | 8881.0 | 2303.0 | 11184.0 |
| 2 | .0 | .0 | 75.0 | .0 | 1600.0 | 3530.0 | 5205.0 |
| 3 | .0 | .0 | .0 | 370.0 | 200.0 | 1814.0 | 2384.0 |
| 4 | .0 | .0 | .0 | .0 | 167.0 | 203.0 | 370.0 |
| 5 | .0 | 5205. | 2309. | .0 | .0 | 3969.0 | 11483.0 |
| 6 | 11184. | .0 | .0 | .0 | 635.0 | .0 | 11819.0 |
| 7 | 11184. | 5205. | 2384. | 370. | 11483.0 | 11819.0 | 42445.0 |

The estimation of the probabilities is now a simple matter of dividing each entry in the 6 × 6 submatrix by the *TST*, which appear in Table 2. The rows and columns in the 6 × 6 submatrix of Table 2 yield the joint probabilities *p*_*ij*_, while the 7^th^ row presents the marginal input probabilities (*p*_*i*_) and the 7^th^ column the marginal output probabilities (*p*_*j*_). The (7,7) entry is now unity, because all probabilities have been normalized by the *TST*.

**Table 2.** Normalized flow and marginal probabilities in Cone Spring.

|  | 1 | 2 | 3 | 4 | 5 | 6 | 7 |
| --- | --- | --- | --- | --- | --- | --- | --- |
| 1 | .0 | .0 | .0 | .0 | .2092 | .0543 | .2635 |
| 2 | .0 | .0 | .0018 | .0 | .0377 | .0832 | .1226 |
| 3 | .0 | .0 | .0000 | .0087 | .0047 | .0427 | .0562 |
| 4 | .0 | .0 | .0000 | .0 | .0039 | .0048 | .0087 |
| 5 | .0 | .1226 | .0544 | .0 | .0 | .0935 | .2705 |
| 6 | .2635 | .0 | .0 | .0 | .0150 | .0000 | .2785 |
| 7 | .2635 | .1226 | .0562 | .0087 | .2705 | .2785 | 1.0000 |

Now it is left to insert these probabilities into Eqn (4) to calculate the contributions of each *p*_*ij*_ to the overall *AMI*, i.e., the *AMI*_*ij*_ (Table 3).

**Table 3.** Contributions of all flows to the AMI (in bits) in rows and columns 1 → 6. Marginal sums appear in row 7 and column 7. Total AMI is in element (7,7).

|  | 1 | 2 | 3 | 4 | 5 | 6 | 7 |
| --- | --- | --- | --- | --- | --- | --- | --- |
| 1 | .0 | .0 | .0 | .0 | .3251 | -.0236 | .3015 |
| 2 | .0 | .0 | -.0035 | .0 | .0069 | .1068 | .1103 |
| 3 | .0 | .0 | .0 | .0362 | -.0080 | .0620 | .0903 |
| 4 | .0 | .0 | .0 | .0 | .0029 | .0047 | .0076 |
| 5 | .0 | .2313 | .1001 | .0 | .0000 | .0292 | .3606 |
| 6 | .4861 | .0 | .0 | .0 | -.0349 | .0 | .4512 |
| 7 | .4861 | .2313 | .0966 | .0362 | .2921 | .1790 | 1.3214 |

The sum of all the 6 × 6 submatrix contributions (*AMI*_*ij*_) yields a value of *AMI* = 1.321 bits, which agrees with the value calculated from Eqn. (2). The marginal contributions of all flows out of the designated row node into other nodes appear in column 7 and those of all flows out of other nodes into the column node are shown in row 7.

As mentioned above, contributions of individual flows may at times be negative, as seen in flows 2→3, 3→5 and 1→6. However, the marginal sums of the *AMI*_*ij*_ always remain non-negative^19^.

With regard to the partition representing *AMI*, Alex Zorach demonstrated how the effective number of trophic levels in a flow network is equal to 2^*AMI*^, whenever the units of *AMI* are bits.^20^ In the case of Cone Spring this turns out to be 2.50, which roughly accords with the visual picture in Figure (2).

As for the contributions of all the flows to the conditional entropy (the *Φ*_*ij*_s), they appear in Table 4 according to Eqn. (5), The marginal sums appear in Row 7 and Col 7. The total sum is *Φ* = 1.748 bits.

**Table 4.** Contributions of each flow (in bits) to the conditional entropy. As in Table 3, marginal sums are collected in rows and column 7. Conditional entropy is in (7,7)

|  | 1 | 2 | 3 | 4 | 5 | 6 | 7 |
| --- | --- | --- | --- | --- | --- | --- | --- |
| 1 | 0 | 0 | 0 | 0 | 0.1472 | 0.2518 | 0.399 |
| 2 | 0 | 0 | 0.0196 | 0 | 0.1714 | 0.1916 | 0.3826 |
| 3 | 0 | 0 | 0 | 0.0234 | 0.444 | 0.1324 | 0.2003 |
| 4 | 0 | 0 | 0 | 0 | 0.0285 | 0.0322 | 0.0607 |
| 5 | 0 | 0.14 | 0.1284 | 0 | 0 | 0.2906 | 0.559 |
| 6 | 0.021 | 0 | 0 | 0 | 0.1256 | 0 | 0.1466 |
| 7 | 0.021 | 0.14 | 0.1481 | 0.0234 | 0.5171 | 0.8986 | 1.748 |

Whence, the full Interaction Capacity is *C = AMI + Φ* = 1.321 + 1.748 = 3.069 bits, which agrees with that calculated from Eqn. (1).

The conditional entropy of a flow network can take on several manifestations. The nature of entropy is most frequently conceived as disorganization – the lack of constraints that provide the system with its organization. Several of these are deleterious to the persistence of the nodes and system, such as decay or heavy environmental stresses, such as temperature and toxins. In flow networks, however, an important ambiguity is the multiplicity of pathways connecting most pairs of taxa. Resources are usually not constrained to follow rigid, unique routes from one given taxon to another. In highly connected networks there are often a multitude of alternative paths that can be followed, so that if one connection is hindered or eliminated, compensatory resources may follow other routes to their destination. Such multiplicity contributes directly to conditional entropy. This plurality can be referred to as “functional redundancy” and is an important mode of repair and survival in ecological and economic communities that is missing from the B-G index of diversity. If too many alternate connections are damaged or extirpated, the system can face collapse and extinction.

### Contributions Particular to Nodes or Flows

Having fully decomposed the B-G Interaction Capacity into its flow-wise components of both *AMI* and *Φ*, one may now examine how much any particular flow or node donates to the partitions. For example, the flow from Detritovores (3) to Carnivores (4) is a single, but important connection that contributes 0.0362 bits to the system *AMI* and 0.00234 bits to the conditional entropy, or 0.0596 bits in all to the Interaction Capacity, C. That 60% of its contribution is to the system *AMI* is understandable, since it is a solitary connection to the bottom half of the system diagram. Nevertheless, it still contributes a full 40% of its activity to the conditional entropy, *Φ*, (almost the complimentary proportion of the total Interaction Capacity), because it participates in 2 of the 5 feedback cycles in the system,^21^ that provide alternative routes for system adjustment. This bias must be balanced by exchanges like Detritus (5) to Detritovores (3), which provides 9% of its activity to *AMI* and 91% to *Φ*.

The extremes in any system are likely to be of interest. The top 5 flow contributions to the AMI are:

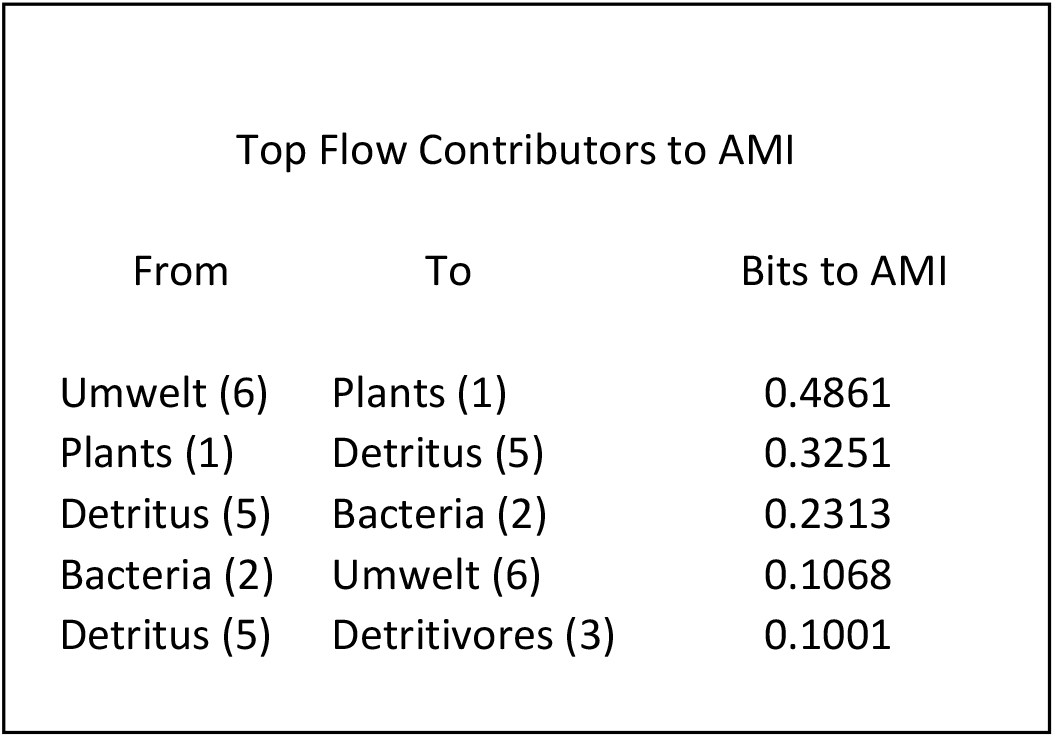

where 6 represents the external world (Umwelt). Together these 5 flows constitute a full 95% of the *AMI*. All are weighted heavily by their flow magnitudes.

Similarly, one can identify the 5 greatest contributors to the conditional entropy as:

| Top Flow Contributors to $\Phi$ | | |
| --- | --- | --- |
| From | To | Bits to $\Phi$ |
| Detritus (5) | Umwelt (6) | 0.2906 |
| Plants(1) | Umwelt (6) | 0.2518 |
| Bacteria (2) | Umwelt (6) | 0.1916 |
| Bacteria (2) | Detritus (5) | 0.1714 |
| Plants(1) | Detritus (5) | 0.1472 |

One notices that the top 5 out of 16 flows can all be identified as conventional dissipative flows. Still, the combined contributions of plants to AMI exceeds the dissipation emanating from them.

It is most fortuitous that the marginal sums of all contributions to either *AMI* or *Φ* remain non-negative (row and column 7 in Table 3.) This allows an investigator to assign the full contributions of each node to the Interaction Capacity (C). (This would be analogous to identifying the contribution of each node to the conventional B-G diversity.) The marginal sums of the contributions to the Interaction Capacity are not symmetrical, however. That is *AMI*_*i*_ is ≠ to *Φ*_*i*_ for all *i*, because the marginal sums of the nodes as donors do not always equal the marginal sums of those counted as receptors. Nevertheless, each flow contribution is counted exactly twice, so that averaging the donor and receptor marginal sums of each node yields symmetry in their averages.

Executing the average nodal marginal contributions for the Cone Spring system yields definable nodal partitions (Table 5):

**Table 5.** Distribution of marginal nodal contributions to AMI and Φ.

| *Taxon* | *AMI<sub>i</sub> (bits)* | *Φ<sub>i</sub> (bits)* | *C<sub>i</sub> (bits)* | *% of C* |
| --- | --- | --- | --- | --- |
| Plants(1) | .3938 | .2010 | .5948 | 19.43 |
| Bacteria(2) | .1708 | .2613 | .4321 | 14.12 |
| Detritivores(3) | .0934 | .1742 | .2676 | 8.74 |
| Carnivores(4) | .0219 | .0421 | .0640 | 2.09 |
| Detritus(5) | .3264 | .5381 | .8645 | 28.25 |
| Umwelt(6) | .3151 | .5226 | .8377 | 27.37 |
| Totals | 1.321 | 1.748 | 3.069 | 100.00 |

It appears that in only one case (Plants, #1) do their contributions to constraints exceed their amendments to flexibility. This reflects their role as a primary driver, providing almost all of the energy and material to the rest of the system.

While the flow network of the Crystal River Marsh Inlet contains 91 separate flows, making reporting the results on a flow basis cumbersome, the marginal contributions of its 18 nodes, however, are conveniently reported here in Table 6, ranked according to their shares of contributions to the Interaction Capacity, C.

**Table 6.**
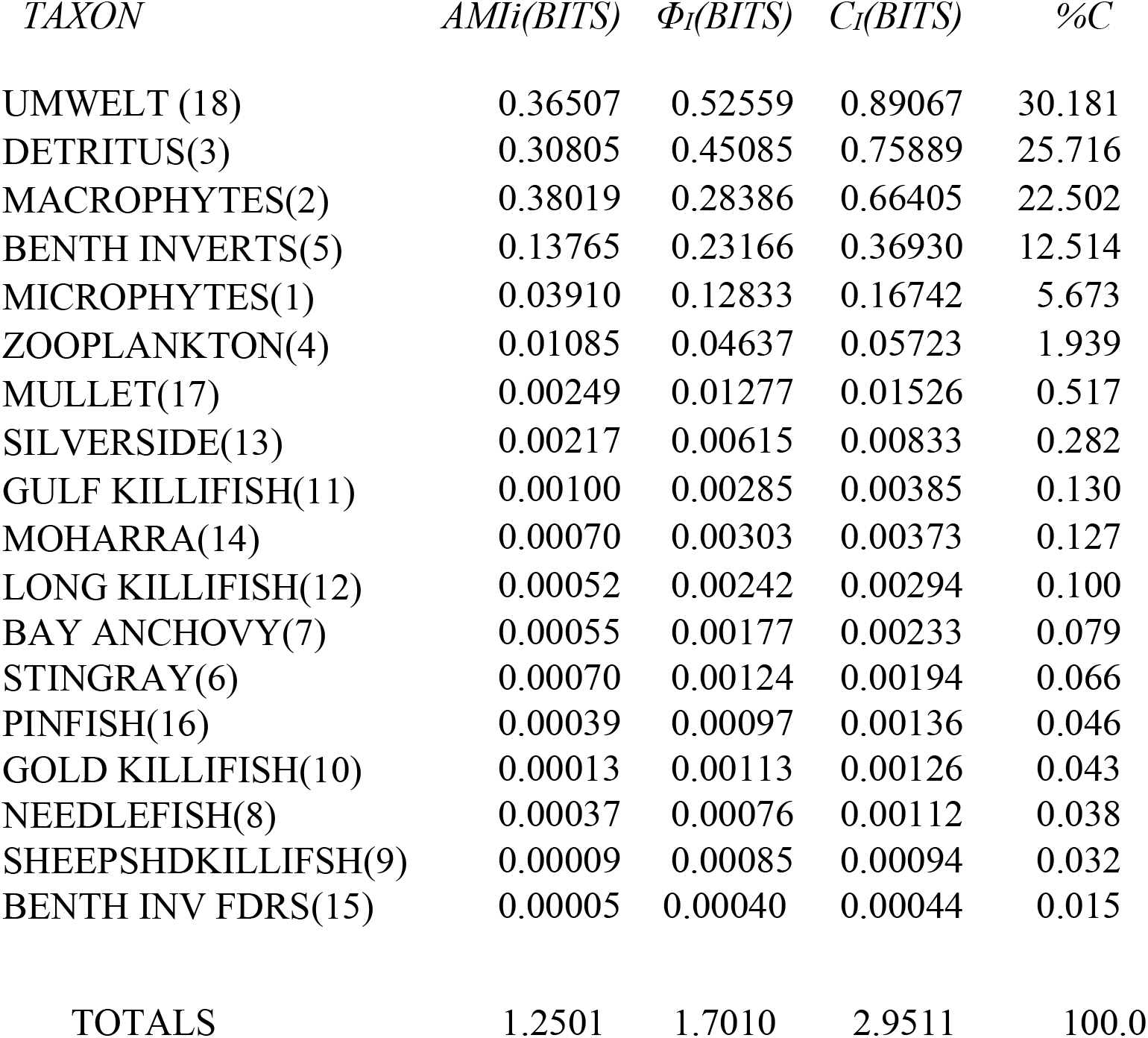
Marginal Contributions of taxa in Crystal River Marsh, ranked by %C.

As with the Cone Spring system, in only one case (Macrophytes) does its contribution to *AMI* exceed its addendum to *Φ*, reflecting its role as the driver of the system.

Again, these various contributions are modulated by the magnitudes of their flows. If, however, one is interested in adjusting the flows and topologies of a network, one seeks to identify those flows that might impart significant change to the ratio of *AMI:Φ* so as to move the system toward the center of the window of vitality^22^. Toward that goal, one would like to calculate the sensitivities of the contributions of the flow elements to changes in their joint probabilities, i.e., *∂AMI/∂p*_*ij*_ and *∂Φ/∂p*_*ij*_.

Fortunately, such indicators can be determined without much further labor. One notes In that regard that the formulae for *AMI* and *Φ* are first-order homogeneous Euler functions of the form, *f(x,y) = x g(x,y)*.^23^ In that case, one may apply Euler’s theorem regarding the derivatives of such functions, namely,

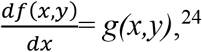

So that 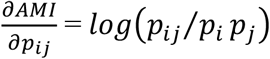, and 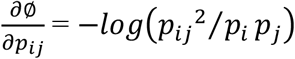. That is, the sensitivities of AMI and Φ are simply the contributions divided by their respective *p*_*ij*_ s. For the Cone Spring network the flow sensitivities are calculated and shown in Table 7 as:

**Table 7.**
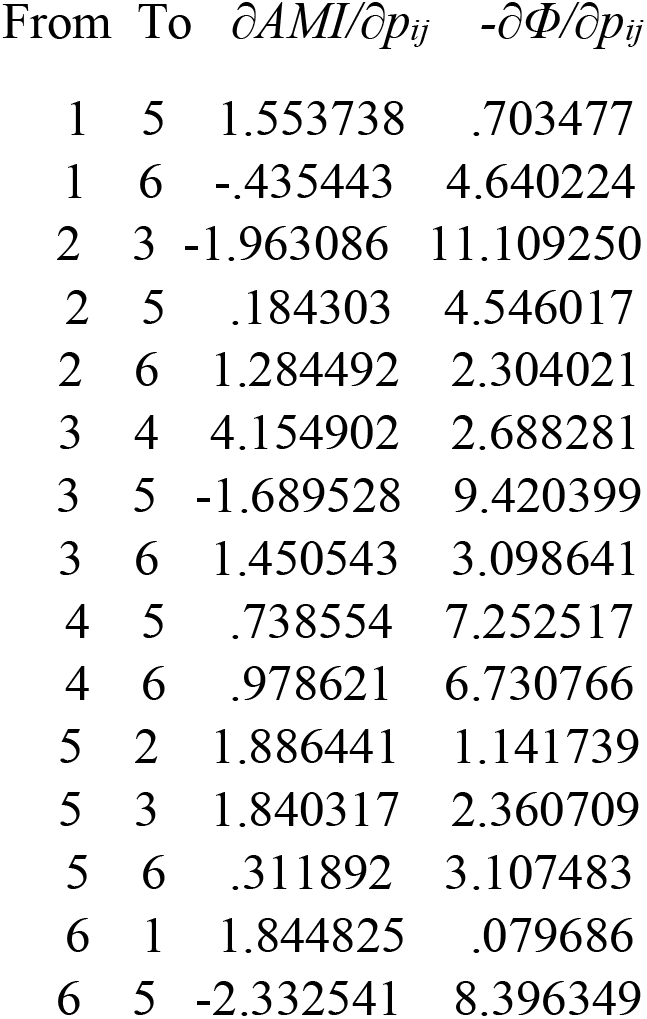
Sensitivities of AMI and Φ to increases in flows between components.

| From | To | $\partial AMI/\partial p_{ij}$ | $-\partial \Phi/\partial p_{ij}$ |
| --- | --- | --- | --- |
| 1 | 5 | 1.553738 | .703477 |
| 1 | 6 | -.435443 | 4.640224 |
| 2 | 3 | -1.963086 | 11.109250 |
| 2 | 5 | .184303 | 4.546017 |
| 2 | 6 | 1.284492 | 2.304021 |
| 3 | 4 | 4.154902 | 2.688281 |
| 3 | 5 | -1.689528 | 9.420399 |
| 3 | 6 | 1.450543 | 3.098641 |
| 4 | 5 | .738554 | 7.252517 |
| 4 | 6 | .978621 | 6.730766 |
| 5 | 2 | 1.886441 | 1.141739 |
| 5 | 3 | 1.840317 | 2.360709 |
| 5 | 6 | .311892 | 3.107483 |
| 6 | 1 | 1.844825 | .079686 |
| 6 | 5 | -2.332541 | 8.396349 |

One immediately notices how increments in flows with negative contributions to AMI seem to cause the greatest relative changes to *AMI* and *Φ*. If these sensitivities seem to be rather large, it is because they represent a unit increase in each *p*_*ij*_, most of which are quite small to begin with. Nonetheless, all sensitivities can serve as indicators of which flows to change and in which direction so as to move the system towards any desired goal.^25^

### Does Current Ecosystem Research Adequately Address Ecoduality?

The foregoing exercises require only estimated data and the simple algebra of logarithms. As such they are phenomenological (any decisive causalities are absent), rather than theoretical, or even philosophical, in nature. Unfortunately, phenomenology is eschewed by most ecologist, who are motivated more to solve problems in the manner of physics. This usually involves mechanical models as tools of simulation, and the models require the supposition of distinct causes. (Let no one denigrate this conventional approach, as it has yielded innumerable insights into ecosystem events, even for the author.)

The unspoken assumption remains, however, that, if enough situations were clarified, a unified theory of ecosystem behavior would emerge from the endeavor. It remains unclear, however, whether such will be the outcome of the ecological project. A while ago, Kirstin Shrader-Frechette asked the author whether, in the end, ecology might resemble just a large collection of case studies?^26^ If, in fact, the well-developed theoretical body of physics does not fully consider *all* classes of phenomena, the body of ecosystem studies could indeed remain unsatisfactorily incomplete. Four such classes that currently are inadequately addressed and deserve more intense examination are: (1) relationships, (2) processes, (3) apophasis and (4) extreme heterogeneity.

The major focii of physical reality are generally objects. Relationships are usually relegated to the boundary-value problem, if considered at all. Objects generally are assumed to take priority over processes, as is the case in thermodynamics, where the latter are considered to originate from objects, while flows or processes remain only of derivative, secondary interest.

Furthermore, as Gregory Bateson pointed out^27^, physics almost completely ignores that which is missing, and the fundamental force laws of physics remain applicable *only* to homogeneous variables.^28^

The nature of ecosystems as treated here, seems to infer that investigation needs to step beyond the limits of physics. But outside that box, many or most causes remain unknown or even unacknowledged. The alternative, then, is to engage in phenomenology – the search for reliable quantitative description, absent any notion of causality. Here two points are worth mentioning: (a) Almost all accepted principles began with phenomenological searches and later evolved into laws. (b) If there is any conflict between theory and phenomenology, the rules of science require that phenomenology take precedence over theoretical hypotheses.

It was to networks of ecological actors (taxa) that a reasonable minority of ecologists has begun to explore and quantify relationships, heterogeneity and eco-processes.^29^ Numerous algorithms have been written to quantify the nature of these properties.^30^ A less popular route has been to invoke the algebra of information theory to quantify missing constraints.^31^

It bears repeating that information theory rests, not upon the information defined by constraints (i.e., those positivist factors that impart order to a system), but rather begins with the *absence* of constraints (uncertainty.) As discussed here, the Capacity of Interactions can be separated into positivist tendencies and apophatic vacancies. Two different types of interdependent causalities govern the two realms – (a) the positivist (and often contingent) forces of nature, and (b) the ever-increasing vacuity that admits room for the positivist propensities to make new structure. The algebra of information theory shows how the two activities are opposing, but interdependent.^32^

Whence it is that Fitzgerald’s “special intelligence” need not be particularly special. In this ecological consideration, the dual functionalities differ in terms of causal origins and their natures. A partial resolution of the seeming paradox appears when one realizes that the contradictory actions of positivist forces and entropic flexibilities predominate at different hierarchical levels of action. The contribution of a flow or node to the *AMI* derives mostly from competition and cooperation among the taxa that builds niches for the participating species.

Augmentation of the conditional entropy, *Φ*, by contrast, involves how the diverse elements support system-level flexibility and thereby some security for the whole system.

The fact that these actions pertain as well to any system that can be represented as a two-dimensional network implies that the dualism generated by ecosystems has its counterparts in numerous other systems, such as social groups, fluid flow networks, electrical grids, traffic street patterns, or supply chains, whether of natural or anthropogenic origin.^33^ Knowing how to quantify *all* contributions provides a valuable guide to system management or design.

While much is yet to be gained in ecology by continuing with material, mechanical models, even broader progress seems to be possible by striving to fulfill Eugene Odum’s valedictory,

*“To achieve a truly holistic or ecosystematic approach, not only ecology, but other disciplines in the natural, social and political sciences as well must emerge to new hitherto unrecognized and unrestricted levels of thinking and action*.*”*^*34*^

Ecologists need to explore pushing harder against the limits of conventional physics.

## Acknowledgements

The author wishes to thank Charles W. Carter, Jr. for suggesting to him that duality is more widespread than just photon materiality and wave action, and to Brian Fath for urging this project on him long ago and providing comments on an early draft of this work.

